# Comparative genomics of *Acinetobacter baumannii* and therapeutic bacteriophages from a patient undergoing phage therapy

**DOI:** 10.1101/2021.12.14.472731

**Authors:** Adriana Hernandez, Mei Liu, James Clark, Tram Le, Biswajit Biswas, Kimberly A. Bishop-Lilly, Matthew Henry, Javier Quinones, Theron Hamilton, Robert Schooley, Scott Salka, Ry Young, Jason Gill

## Abstract

In 2016, a 68-year-old patient with a disseminated multi-drug resistant *Acinetobacter baumannii* infection was treated using lytic bacteriophages in one of the first modern human clinical uses of phage therapy in the United States. Due to the emergency nature of the treatment there was little time to thoroughly characterize the phages used in this intervention or the pathogen itself. Here we report the genomes of the nine phages used for treatment and three strains of *A. baumannii* isolated prior to and during treatment. The eight phages used in the initial treatment were found to be a group of closely related T4-like myophages; the ninth phage, AbTP3Φ1, was found to be an unrelated Fri1-like podophage. Analysis of 19 *A. baumannii* isolates collected before and during phage treatment showed that resistance to the T4-like phages appeared as early as two days following the start of treatment. Three *A. baumannii* strains (TP1, TP2 and TP3) collected before and during treatment were sequenced to closure, and all contained a 3.9 Mb chromosome of sequence type 570 with a KL116 capsule locus and identical 8.7 kb plasmids. Phage-insensitive mutants of *A. baumannii* strain TP1 were generated *in vitro* and the majority of identified mutations were located in the bacterial capsule locus. The presence of the same mutation in both the *in vitro* mutants and in phage-insensitive isolates TP2 and TP3, which evolved *in vivo* during phage treatment, indicate that *in vitro* investigations can produce results that are relevant and predictive for the *in vivo* environment.

## Introduction

The Gram-negative bacterium *Acinetobacter baumannii* is recognized as one of the most important pathogens in healthcare-associated infections, particularly with ventilator-associated pneumonia and catheter associated infections (Dijkshoorn, Nemec et al. 2007, Peleg, Seifert et al. 2008, Lee, Lee et al. 2017). This is especially true for carbapenem-resistant *A. baumannii*, which caused 8,500 infections and 700 deaths in the U.S. in 2017 alone (CDC 2019). Several characteristics of this pathogen impact treatment regimens and outcomes, including the increased prevalence of multidrug-resistant (MDR) strains, environmental persistence due to its desiccation and disinfectant resistance, biofilm formation, and motility (Roca, Espinal et al. 2012, Harding, Hennon et al. 2018). This results in hampered clinical intervention strategies and increased risks of reinfection and outbreaks (Chusri, Chongsuvivatwong et al. 2014). As cases of resistant infections are more prevalent and very few new antibiotics are available, the use of bacteriophages (phages) to treat and/or control multidrug-resistant infections is being reconsidered as an alternative strategy for therapeutic and prophylactic applications (Young and Gill 2015, Nobrega, Vlot et al. 2018, Gordillo Altamirano and Barr 2019)

In the modern era, the first published emergency intervention using phage in treating a systemic multi-drug-resistant *A. baumannii* infection in the US was the well-publicized “Patterson case” in 2016 (Schooley, Biswas et al. 2017). Clinical interventions using phage therapy to combat MDR bacterial infections have increased significantly in the past several years, with successful phage treatment outcomes reported in a number of case studies involving MDR *Pseudomonas aeruginosa, Staphylococcus aureus*, and *Escherichia coli* (Aslam, Lampley et al. 2020). These case studies have been encouraging in terms of clinical outcome, but in-depth examination of the phage-host interaction during treatment and their implications for phage efficacy remains an area of active study.

In principle, the effectiveness of the phage treatment depends on the ability of phage to localize to and persist in the infected tissue and propagate lytically. During this process, both the phages and their bacterial hosts replicate and evolve, potentially reducing the ability of the phages to clear the infection. In the 2016 *A. baumannii* clinical intervention, emergence of phage resistance was reported 8 days following the initiation of phage treatment (Schooley, Biswas et al. 2017). Due to the rapid response required for the 2016 clinical intervention, both the *A. baumannii* pathogen and the phages used in treatment were largely uncharacterized. Here we examine the genomics of the therapeutic phages, the emergence of phage resistance during treatment, and the *in vivo* evolution of the pathogen with complete genomes of three *A. baumannii* strains isolated before and during phage therapy. Genetic changes responsible for phage resistance developed *in vivo* are compared to resistance developed in *vitro*, and the implications for optimizing phage therapeutic interventions are discussed.

## Materials and Methods

### A. baumannii clinical isolates

As reported previously (Schooley, Biswas et al. 2017), *A. baumannii* clinical isolates were isolated from multiple drains, peritoneal fluid, and respiratory secretions of the patient receiving phage treatment at the UCSD Clinical Microbiology Laboratory. Strain TP1 was isolated from peritoneal drain on Feb 10, 2016, strain TP2 and TP3 were isolated from a pancreatic drain on March 21 and March 23, 2016, respectively. All *Acinetobacter* strains were routinely cultured on tryptic soy broth (TSB, 17g/L Bacto tryptone, 3 g/L soytone, 2.5 g/L D-glucose, 5 g/L NaCl, 2.5 g/L disodium phosphate) or Tryptic Soy Agar (TSB plus 1.5% Bacto agar, w/v). For all plaque assays, a 0.5% TB agar overlay (10 g/L tryptone, 5 g/L NaCl and 0.5% Bacto agar) was inoculated with 0.1 ml of a fresh overnight TSB culture of host and poured over TSA plates. All strains were grown at 37 °C.

### Phage propagation, whole genome sequencing and characterization

Except for AB-Navy71, the isolation and propagation of all phages used in three cocktails, ΦPC, ΦIV, and ΦIVB were conducted using the soft agar overlay method (Adams 1959), and were described in detail previously (Schooley, Biswas et al. 2017). Phage AB-Navy71was purchased from the Leibniz Institute DSMZ (www.dsmz.de) as phage name vB-GEC_Ab-M-G7 (DMS25639). Phage DNA was extracted using the Promega Wizard DNA extraction system following a modified protocol as previously described (Summer 2009). The DNA was prepared for sequencing with 550 bp inserts using a TruSeq Nano kit and sequenced as paired end 250 bp reads by Illumina MiSeq with V2 500-cycle chemistry. Reads were checked for quality using FastQC (www.bioinformatics.babraham.ac.uk/projects/fastqc) and the genome was assembled using SPAdes v3.5.0 (Bankevich, Nurk et al. 2012). The assembled contigs were completed by running PCR amplifying the region covering the contig ends, sequencing the resulting PCR products (see Supplementary Table S1 for PCR primers used), followed by manual verification. Annotation of the assembled genome was conducted using tools in Galaxy hosted by https://cpt.tamu.edu/galaxy-pub (Afgan, Baker et al. 2018). Genes were identified using Glimmer v3 (Delcher, Harmon et al. 1999) and MetaGeneAnnotator v1.0 (Noguchi, Taniguchi et al. 2008), and tRNAs were identified using ARAGORN v2.36 (Laslett and Canback 2004). The identified genes were assigned putative functions using default settings of BLAST v2.9.0 against the nr and SwissProt databases (Camacho, Coulouris et al. 2009, UniProt Consortium 2018), InterProScan v5.33 (Jones, Binns et al. 2014), and TMHMM v2.0 (Krogh, Larsson et al. 2001). For comparative purposes, whole genome DNA sequence similarity was conducted using ProgressiveMauve v2.4 (Darling, Mau et al. 2010). Genome maps were made using the linear genome plot tool, and genome comparison maps were made using X-vis, a custom XMFA visualization tool developed by the CPT. Phylogenetic tree of the phage tail fiber proteins was constructed by aligning the protein sequences with MUSCLE (Edgar 2004), and using the pipeline available at https://www.phylogeny.fr/ (Dereeper, Guignon et al. 2008) to run the maximum likelihood analysis (Anisimova and Gascuel 2006). The tree was plotted using TreeDyn (Chevenet, Brun et al. 2006). Tail fiber protein multiple sequence alignment was illustrated using Clustal Omega under default settings (Madeira, Park et al. 2019). Except web-based analysis, most analyses were conducted via the CPT Galaxy and WebApollo interfaces (Dunn, Unni et al. 2019, Jalili, Afgan et al. 2020, Ramsey, Rasche et al. 2020) under default settings (https://cpt.tamu.edu/galaxy-pub).

### Determination of phage sensitivity on clinical strains

Phage sensitivity of *A. baumannii* clinical isolates was determined by spotting serially diluted phage suspensions onto bacterial lawns produced by the soft agar overlay method (Adams 1959). Aliquots of 10 μl serially diluted phage were spotted onto the agar overlay plate, which was incubated at 37°C for 18-24 h to observe plaque formation. All assays were performed in triplicate.

### Phenotype microarrays

Omnilog Phenotype Microarray panels 1-20 for bacterial strains and Dye Mix D (Biolog; Hayward, CA) were used for phenotypic profiling of pancreatic drainage isolates TP1 and TP3. Each strain was assayed with three independent replicates for 48 hours following manufacturer’s instructions. Area under the curve (AUC) values were analyzed using the R package opm (Vaas, Sikorski et al. 2013).

### Genome sequencing and genome analysis of *A. baumannii* TP1, TP2, and TP3

Genomic DNA of *A. baumannii* TP1, TP2, and TP3 was extracted using a bacterial genomic DNA extraction kit from Zymo Research. The extracted DNA was sequenced via Illumina TruSeq in parallel to Oxford Nanopore MinIon R9 flowcell sequencing conducted at Texas A&M Institute for Genome Sciences and Society (TIGGS) located in College Station, TX. For Illumina sequencing, libraries were prepared using TruSeq Nano kit and sequenced by Illumina MiSeq with V2 500-cycle cartridge. For Oxford Nanopore MinIon R9 flowcell sequencing, a Nanopre SQK-RAD004 Rapid Sequencing Kit was used. Using reads obtained from Illumina and Nanopore, complete genome sequences of TP1, TP2, and TP3 were determined using a combination of bioinformatic tools and conducting confirmational PCR for gap regions (see Supplementary Table S1 for PCR primers used). Illumina reads were passed through FastQ groomer (Blankenberg, Gordon et al. 2010) and trimmed using Trimmomatic (Bolger, Lohse et al. 2014) with parameter settings AVGQUAL= 25; SLIDINGWINDOW = 4, average quality required = 28; TRAILING = 25. After trimming, reads were checked for quality with FastQC (www.bioinformatics.babraham.ac.uk/projects/fastqc). For TP1, Nanopore reads were trimmed, and initial assembly was performed with Unicycler to generate a scaffolding draft genome. For TP2 and TP3, trimming was not performed and Canu was used to create a *de novo* assembly using raw reads. Illumina reads were mapped onto the draft genomes with Bowtie2 (Langmead, Trapnell et al. 2009, Langmead and Salzberg 2012), using fast end-to-end parameters and default settings. These mapped reads were used as input for Pilon (Walker, Abeel et al. 2014) under default settings with variant calling mode OFF. After initial sequence corrections the Illumina reads were mapped to the updated genome with Bowtie2 set to sensitive end-to-end mapping. Pilon was run again with newly mapped reads to produce the final output. All other settings remained default. All reads were then remapped against the Pilon-produced contig, and low coverage areas or areas with ambiguous base calls were confirmed by PCR (see Supplementary Table S1 for PCR primers used). The complete genome sequences were deposited to NCBI and annotated by the NCBI Prokaryotic Genome Annotation Pipeline (PGAP) (Tatusova, DiCuccio et al. 2016). The closed, circular genome sequences were re-opened upstream of *dnaA*.

Antibiotic resistance genes (ARGs) were identified using the CARD Resistance Gene Identifier (https://card.mcmaster.ca/) allowing for perfect and strict hits (Alcock, Raphenya et al. 2020) under default settings. The capsule (K) locus was identified using the Kaptive web interface (Wick, Heinz et al. 2018, Wyres, Cahill et al. 2020). Prophage regions were detected using PHASTER (Arndt, Grant et al. 2016) and the boundaries verified by BLASTn against related bacterial genomes and identification of *attL* and *attR* sites as direct repeats. Through a workflow developed at the CPT (https://cpt.tamu.edu/galaxy-pub), the prophage regions were compared to phage and bacterial genomes available in the NCBI nt database via BLASTn (Camacho, Coulouris et al. 2009), and ProgressiveMauve (Darling, Mau et al. 2010) was used to calculate percent identities. The location and size of indels and SNPs in TP2 and TP3 in reference to TP1 were determined by using ProgressiveMauve (Darling, Mau et al. 2010), followed by manual verification.

### Generation of phage-resistant *A. baumannii* mutants *in vitro*

Phage-resistant mutants of *A. baumannii* TP1 were generated *in vitro* by spotting undiluted phage lysates (10 μl) to lawns of TP1 and picking colonies growing within the spots following overnight incubation, and streaking to fresh TSA plates. These isolates were then used to inoculate fresh TSB cultures which were grown to an OD_550_ of 0.2 - 0.3 and infected with the same phage at an MOI of 0.2. The cultures were incubated for 6 hours at 37 °C, and then plated on TSA to produce individual colonies. A single colony was isolated from these plates and purified by an additional round of subculture. Strains were confirmed to be resistant to phage by spot assays as described above. Three independent phage-resistant mutants were isolated against phages AC4, Maestro, AB-Navy97, AbTP3phi1, and two independent mutants were isolated against phage AB-Navy1.

### Genomic characterization of *in vitro* phage-resistant *A. baumannii* mutants

Genomic DNA of *A. baumannii* was extracted as described above, prepared for sequencing with an Illumina TruSeq Nano kit, and sequenced by Illumina MiSeq V2 for 500 cycles. Bowtie2 (v. 2.2.4) (Langmead, Trapnell et al. 2009) was used to map forward and reverse raw reads to the reference genome of the parental strain TP1 in --fast mode with maximum fragment length set to 800. BAM files were analyzed in samtools mpileup v.1.2 with max per-file depth of 250. Bcftools call v.1.3.0 was used to identify SNPs and indels by consensus call in haploid mode. Read mapping of the parental (TP1) reads against the reference genome was used to subtract spurious variant calls from mapped mutant reads, and remaining variant calls were filtered to retain calls with quality scores of 100 or greater.

### NCBI accession numbers

The genomes of *A. baumannii* TP1, TP2, and TP3 were deposited in the NCBI database under BioProject <u>PRJNA641163</u>, with the following accession and BioSample numbers. TP1: <u>CP056784</u> and <u>SAMN15344688</u>; TP2: <u>CP060011</u> and <u>SAMN15735522</u>; TP3: <u>CP060013</u> and <u>SAMN15738014</u>. Phages were deposited to NCBI under the following accession numbers: <u>MT949699</u> (Maestro), OL770258 (AB-Navy1), OL770259 (AB-Navy4), OL770260 (AB-Navy71), OL770261 (AB-Navy97), OL770262 (AC4), OL770263 (AbTP3Phi1).

## Results and Discussion

### Genomic characterization of phages used in human clinical intervention

The clinical course of the *A. baumannii* infection and phage treatment, known as the “Patterson Case”, has been described previously (Schooley, Biswas et al. 2017). Briefly, phage treatment was initiated with two phage cocktails, each containing four phages: cocktail ΦPC was administered into abdominal abscess cavities through existing percutaneous drains, and cocktail ΦIV was administered intravenously. Near the end of patient treatment, a ninth phage, AbTP3Φ1, was isolated to target the phage-resistant *A. baumannii* strain TP3 that arose during treatment. Phage AbTP3Φ1 was administered intravenously in a two-phage cocktail (phiIVB) in combination with one phage from cocktail phiIV (Schooley, Biswas et al. 2017). As a follow up study to this phage intervention case, we determined the genomes of the phages and also of the bacterial strains that were isolated during phage treatment. All nine phages used in these cocktails were sequenced and their genomes are summarized in **Table 1**. Genome sequences of phages C2P12, C2P21 and C2P24 used in cocktail ΦPC were determined to be identical, so phage C2P24 was renamed as phage Maestro and is used as a representative of this group. The phages can be categorized into two broad groups: phages Maestro, AC4, ABphi1, ABphi4, ABphi71 and ABphi97 are large (165-169 kb) T4-like myophages, and phage AbTP3Φ1 is a 42 kb podophage.

The six myophages used are closely related, with nucleotide sequence identity ranging from 64.2%-95.6% between any two genomes, as determined by progressiveMauve analysis (**Figure 1**). Based on analysis by BLASTn, both Maestro and AB-Navy71 share 90%-96% overall sequence identity with *Acinetobacter* phage AbTZA1 (NC_049445), which is classified as a member of the genus *Hadassahvirus* by the International Committee on Taxonomy of Viruses (ICTV) (Adriaenssens, Krupovic et al. 2017); predicted taxonomic placements of the other four closely related myophages (AC4, AB-Navy1, AB-Navy4, AB-Navy97) are in the genus *Lazarusvirus* based on 92%-96% sequence identity shared between each phage with phage AM101 (NC_049511).

**Figure 1.**
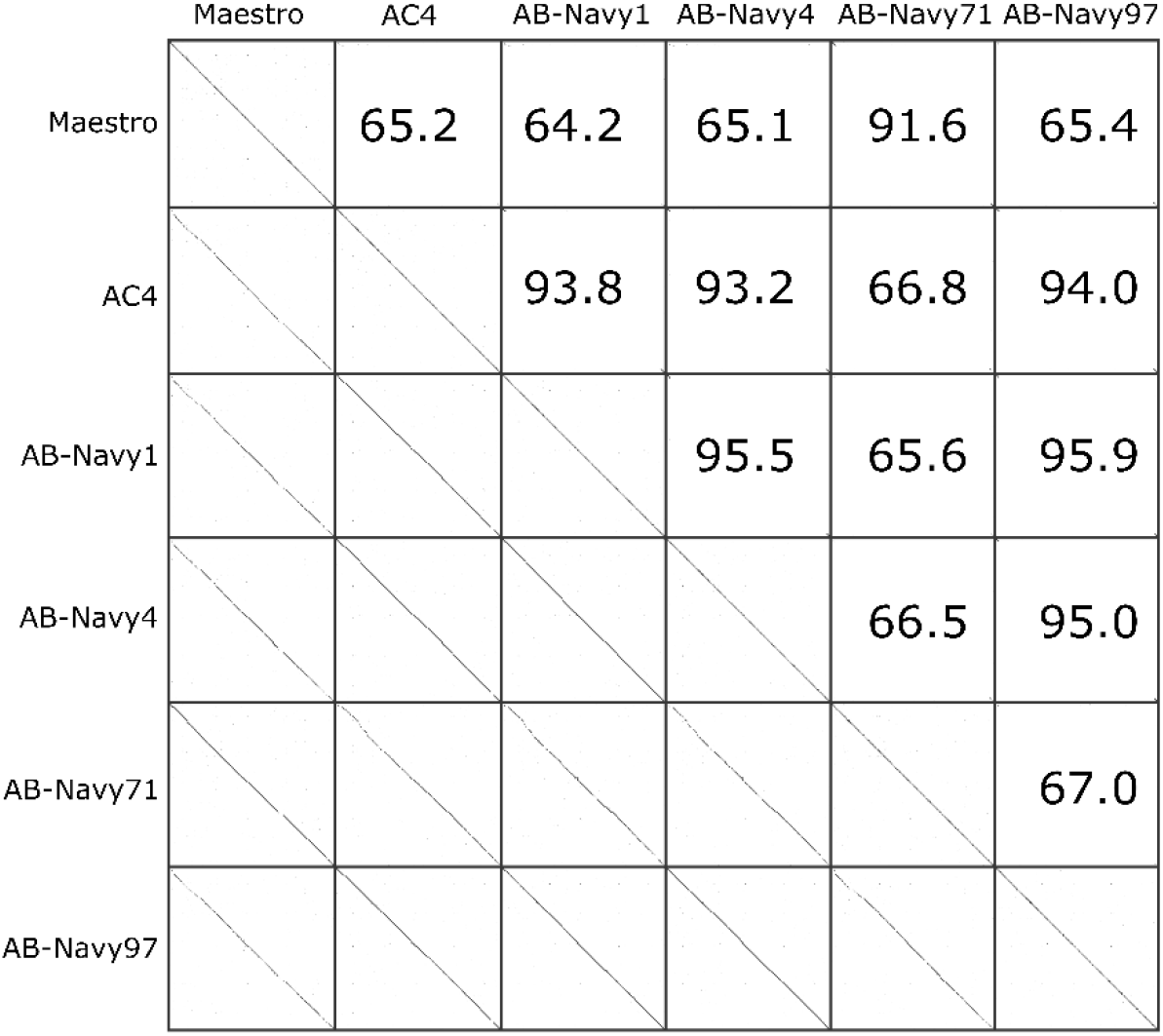
DNA sequence relatedness of six T4-like phages. Upper section: pairwise percent DNA sequence identities between all six phages, as determined by ProgressiveMauve. Lower section: dotplots visually representing DNA sequence homology between phages.

The genome of Maestro is presented as an example for this group of *Acinetobacter* myophages (**Supplementary Figure 1**). Maestro has a complete genome size of 169,176 bp and a GC-content of 36.6%. Seven tRNA genes were identified, including one that appears to specify an amber codon. Genes encoding phage integrases or proteins associated with bacterial virulence were not detected. A conserved core of 95 T4-like genes were identified, clustered in several regions of the genome. These include genes encoding structural proteins and proteins involved in DNA nucleotide metabolism and replication. Proteins involved in transcriptional regulation in T4 were found to have homologs in Maestro, which suggests Maestro follows a T4-like program of gene expression, with positive control of early, middle and late transcripts (Miller, Kutter et al. 2003). Interspersed between conserved gene clusters are hypothetical ORFs with no clear associated function, mostly conserved among T4-like phages infecting *Acinetobacter* but not with T4, suggesting that these genes might be involved in host-specific phage interactions. Conserved hypothetical proteins among these T4-like phages of *Acinetobacter* represent 42% of the ORFs in the Maestro genome. The holin and endolysin lysis genes in Maestro are similarly located as in T4 and have high primary structure similarity, indicating that the first two steps in lysis, the permeabilization of the inner membrane and the degradation of the cell wall are effected the same way (Cahill and Young 2019). The third step, disruption of the outer membrane, is accomplished in most dsDNA phages by spanin proteins, encoded by the *pseT.3* and *pseT.2* genes in T4. Even in large genomes like Maestro, spanin genes are detectable because every spanin complex has at least one OM lipoprotein, thus requiring a lipobox motif in the N-terminal amino acid sequence (Kongari, Rajaure et al. 2018). No suitable lipobox motifs were detected in the Maestro genome, indicating that Maestro, like some other *Acinetobacter* phages, uses a different mechanism for OM disruption (Hernandez-Morales, Lessor et al. 2018, Kongari, Rajaure et al. 2018). Homologs of the phage T4 RI and RIII antiholins were identified in the Maestro genome, indicating this phage has the ability to undergo T4-like lysis inhibition (Krieger, Kuznetsov et al. 2020). The effects of lysis inhibition in therapeutic interventions are not known, but superinfection-induced lysis inhibition delays lysis time and increases burst size *in vitro* and could affect *in vivo* phage proliferation at the site of therapeutic application.

During the infection process of phage T4, the long tail fibers (LTFs) bind to the phage’s receptor on the cell surface. In T4, the LTF is comprised of Gp34, Gp35, Gp36 and Gp37, which form the proximal LTF, two joints, and distal LTF, respectively; the distal LTF contains the phage receptor-binding function in its C-terminal domain (Bartual, Otero et al. 2010, Hyman and van Raaij 2018). The LTFs and receptor-binding proteins of the myophages used in phage treatment were identified based on their similarity to T4 proteins. The distal domains of the myophage LTFs, containing the predicted receptor binding domains, were compared by multiple sequence alignment (**Supplementary Figure 3**) and construction of a neighbor-joining tree to determine their relationships (**Figure 2**). Multiple sequence alignment revealed the myophages used in the cocktails had two different types of tail fibers, with Maestro, AC4, Navy71 belonging to one cluster, and Navy1, Navy4, and Navy97 belong to the other cluster (**Figure 2**). This finding correlates with the phage resistance patterns observed in *A. baumannii* strains isolated from the patient before and during phage treatment (**Table 2**). Strains resistant to phage AC4 were also resistant to phage Maestro as well as to AB-Navy71, but the same strains were still partially sensitive to Navy1, Navy4, and Navy97. Six days after the start of treatment, resistance to phage AB-Navy1, AB-Navy4 and AB-Navy97 was observed simultaneously.

**Figure 2.**
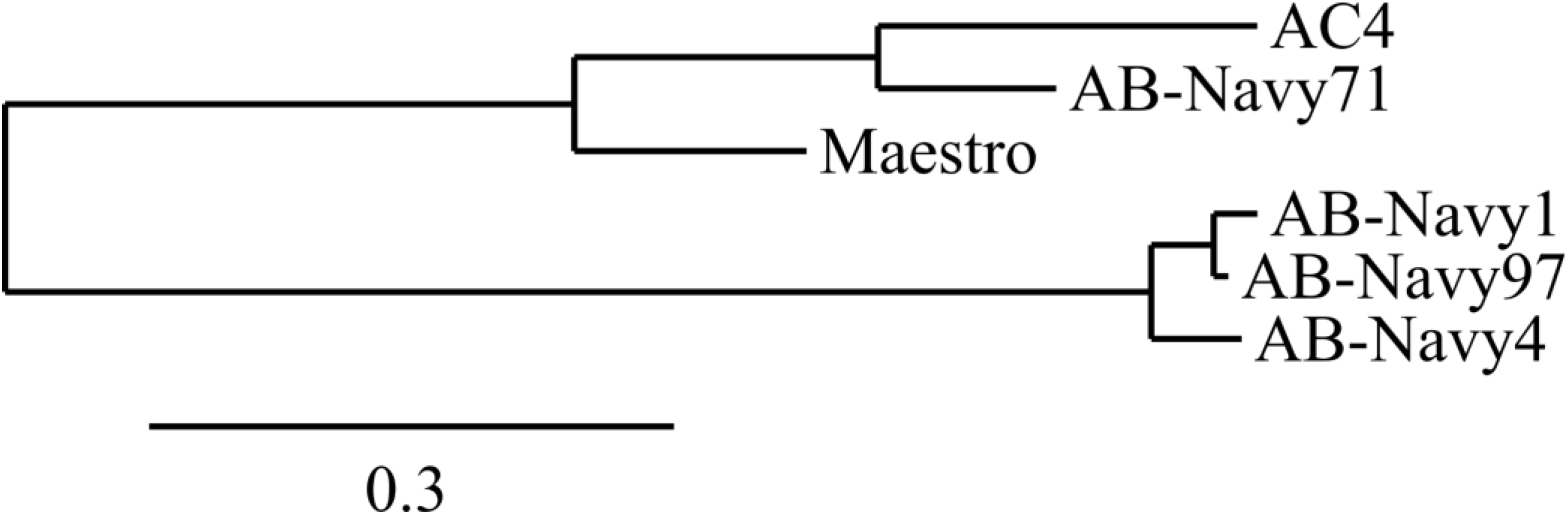
Phylogenetic tree constructed based on the long tail fiber protein sequences of the myophages used in phage treatment.

The podophage AbTP3Φ1 was not available until near the end of treatment, but this phage is genetically distinct from the myophages and appears to use a different receptor, as *A. baumannii* isolates that are resistant to the myophages appear to retain at least partial sensitivity to this phage. Compared to the myophages described above, the much smaller 42 kb podophage AbTP3Φ1 is classified as a member of the genus *Friunavirus* of the *Autographivirinae* family by ICTV (Adriaenssens, Krupovic et al. 2017) (**Table 1**). The genome map of AbTP3Φ1 is shown in **Supplementary Figure 2**. It shares at 82-89% overall DNA identity, as well as genome synteny, to a group of previously described *Acinetobacter* phages, including IME200 (NC_028987), vB_AbaP_AS11 (NC_041915), Fri1 (KR149290) (Popova, Lavysh et al. 2017) and Pipo (MW366783). As a conserved feature of this group of phages, a terminal repeat region of 396 bp was identified in AbTP3Φ1 genome by the PhageTerm tool (Garneau, Depardieu et al. 2017). The tail spike protein of AbTP3Φ1 shares ≥ 95% identity to those found in this group of phages based on BLASTp alignment, and HHpred searches indicate its tailspike matches the phiAB6 tail spike (5JSD, 99.93%), indicating that AbTP3Φ1 adsorption is associated with exopolysaccharide degradation (Lee, Tu et al. 2017). Similar to the myophages reported in this study, spanin proteins were not found in the genome of AbTP3Φ1 nor in any other *A. baumannii* podophage genomes (Hernandez-Morales, Lessor et al. 2018). Recently, another type of OM disruption protein, the disruptin, was identified in coliphage phiKT (Holt, Cahill et al. 2021). However, no proteins with sequence similarity to the phiKT disruptin is detectable in the genomes of any of these cocktail phages. It is possible that entirely novel OM disruption proteins are encoded in these cocktail phages (Hernandez-Morales, Lessor et al. 2018).

These phage sequencing results highlight the importance of thorough genomic analysis of phages prior to phage treatment in order to maximize treatment success and minimize effort and consumption of resources. While none of the phages used contain any detectable deleterious genes and appear to be strictly virulent, three of the phages used in cocktail ΦPC were found to be genetically identical. Due to the time constraints imposed by the emergency nature of the clinical intervention, phages C2P12, C2P21 and C2P24 were isolated from environmental samples mere days before their production and administration to the patient, which did not allow for extensive characterization. The other phages used in the initial ΦPC and ΦIV cocktails, while not identical, are closely related and fall into only two groups based on tail fiber similarity. This explains why *A. baumannii* isolates collected as soon as two days after the start of phage treatment were either completely insensitive or markedly less sensitive to all of the myophages used in the initial two cocktails. The initial treatment in this case could have been conducted by a cocktail of only two of the myophages, or perhaps even a single myophage, and plausibly produced a similar outcome. It is difficult to speculate on the role of AbTPSΦ1 in the treatment outcome, as this phage was not introduced until the end of treatment after the patient had already made considerable progress towards recovery. However, if AbTP3Φ1 had been available at the start of treatment, a rational design in the phage cocktail would indicate its inclusion due to its lack of relationship to the other phages and apparent use of a genetically independent receptor. Ideal phage cocktails should not only contain phages possessing an exclusively lytic life cycle and be free of deleterious genes, but should also exhibit genetically independent mechanisms of host resistance. Ideally this would be manifest in the use of different receptors, which may be revealed by thorough characterization before their use as therapeutics.

### Phage and antibiotic sensitivity of *A. baumannii* strains isolated during treatment

During phage treatment, *A. baumannii* isolates were collected from the patient via various drains or bronchial washes. These strains were tested for their phage sensitivity via plaque assays. These showed that as early as 2 days after phage administration, the efficiency of all the phages in the first two cocktails (ΦPC and ΦIV) was reduced when tested against the bacterial strains isolated during treatment, evident by the decreased titers on those strains compared to the initial titers observed with TP1 (**Table 2**). In some cases, only a zone of clearing (but no individual plaques) was observed on the plates at high phage concentrations. Consistent with the myophage tail fiber protein sequence alignment (**Figure 2**), host resistance to phages appeared earlier with Maestro, AC4, and Navy71 as a group, and later with phages Navy1, Navy4, and Navy97 as a group. In comparison, resistance to phage AbTP3Φ01 was not observed in bacterial isolates collected throughout two months of phage treatment, although plating efficiencies of AbTP3Φ01 varied by up to three orders of magnitude on strains collected during treatment (**Table 2**). The emergence of phage resistance early in phage treatment again illustrates the potential benefits of well-characterized and rationally designed phage cocktails in treatment, which could be designed to mitigate the emergence of resistance. It also raises questions on the benefits of continued phage treatment beyond the first ~9 days, since all isolates collected after this time are fully resistant to the phage. While it is possible that the prolonged period of phage administration (over 60 days) was not required to produce the observed clinical outcome, other studies have shown that phage-insensitive mutants of *A. baumannii* exhibit reduced virulence (Regeimbal, Jacobs et al. 2016). Thus, maintaining selection pressure for the phage-resistant phenotype may provide a benefit to continued treatment even after the pathogen has developed phage resistance.

Some strains isolated throughout phage treatment were also tested for their antibiotic resistance profiles by traditional microtiter MIC (**Supplementary Table S2**). In general, the antibiotic resistance profiles of all strains isolated during the course of phage therapy remained consistent, indicating that phage therapy did not have a major impact on antibiotic resistance of the pathogen. Although an initial report indicated resistance to colistin and tigecycline prior to the start of phage therapy (Schooley, Biswas et al. 2017), sensitivity to colistin and tigecycline (in the range of 2-8 ug/ml) was observed in strains isolated ~7 weeks after the start of phage therapy (collected on May 9, 2016). While sensitive to colistin and tigecycline, these strains were resistant to minocycline. We previously reported on a potential synergistic *in vitro* activity between phage cocktail and minocycline (used at sub-inhibitory concentrations of 0.25 ug/ml) in inhibiting bacterial growth (Schooley, Biswas et al. 2017). However, such results were obtained using strain TP3, and TP3 was not tested for its sensitivity to minocycline, colistin, or tigecycline in these MIC assays. The effect of phages on the antibiotic resistance of *A. baumannii* warrants further study.

To more fully delineate the phenotypic differences between TP1 and TP3, BioLog phenotypic microarray profiling was conducted using phenotypic microarrays (PM) 1-20 (**Figure 3, Supplementary Table S3**). As expected given the clonal nature of the isolates, the phenotypic microarrays demonstrated very consistent phenotypes in terms of carbon, nitrogen, phosphorus and sulfur utilization; biosynthetic pathways and nutrient stimulation; osmotic/ionic response; and pH response; as well as very consistent phenotypes in the chemical sensitivity assays (**Figure 3**). The phenotypic profiling results show that growth of both isolates TP1 and TP3 could be inhibited by colistin or minocycline at higher concentrations (**Figure 3**, yellow box and light blue box, respectively); tigecycline sensitivity is not included in the phenotype microarray panel. Isolate TP3 was found to be completely resistant to nafcillin in this assay, whereas TP1 was sensitive (**Figure 3**, purple box).

**Figure 3.**
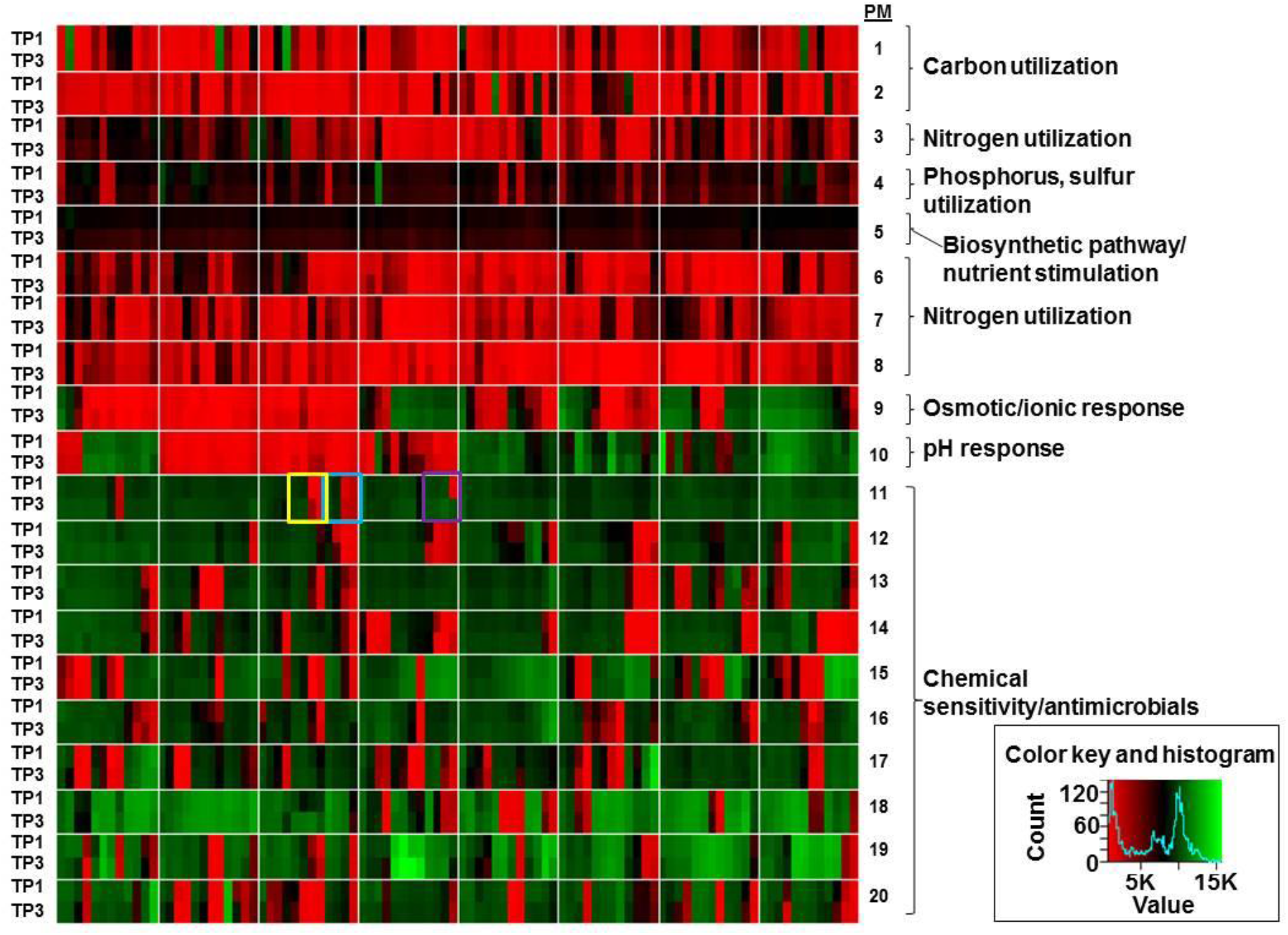
Phenotypic profiling of strains TP1 and TP3. Yellow box: colistin. Light blue box: minocycline. Purple box: nafcillin. Results are calculated using the using area under the curve for 48 hours of growth and are represented are the average of three replicates per strain.

### Characterization of *A. baumannii* strains TP1, TP2, and TP3 isolated before and during phage therapy

Three *A. baumannii* isolates, TP1, TP2, and TP3, were sequenced to closure using a combination of short-read (Illumina) and long-read (Nanopore) sequencing to investigate pathogen evolution during the course of phage treatment. Sequencing to closure allows for comparison of not only SNPs and indels during bacterial evolution but also for tracking of the number and position of mobile DNA elements that are often not assembled into larger contigs if the genomes are only sequenced to a draft state with short-read sequencing. Strain TP1 was isolated prior to the start of phage treatment and was the clinical isolate used to determine phage sensitivity and conduct environmental phage hunts for assembly of therapeutic phage cocktails (Schooley, Biswas et al. 2017). Strains TP2 and TP3 were isolated 6 days and 8 days after the start of phage treatment. All three strains were found to contain a single 3.9 Mb chromosome and a single 8.7 kb plasmid (**Table 3**). Some variation was observed in bacterial chromosome length between strains but the plasmids contained in each strain were identical, and it is clear that these three isolates represent the evolution of strains from a common ancestor over time rather than a succession invasion by different strains. Analysis of the genomes in pubMLST (Jolley, Bray et al. 2018) identified all three isolates as sequence type 570 (Pasteur) and analysis in Kaptive (Wick, Heinz et al. 2018) identified a 20.5 kb region (base position 3,774,031 - 3,794,556 in the TP1 genome) containing 17 genes encoding a predicted capsule type of KL116 (**Supplementary Table S4**) (**Figure 5A**). Consistent with the broad antibiotic resistance observed in these isolates, 32 (TP1) and 35 (TP2, TP3) antibiotic resistance genes were identified based on searches against the CARD 2021 database (Alcock, Raphenya et al. 2020) (**Supplementary Table S5**). The 8.7 kb plasmid contained in TP1, TP2 and TP3 does not encode any identifiable AMR genes, and is identical to plasmids carried in many other *A. baumannii* strains deposited in NCBI. Less than 200 SNPs and indels were observed between these isolates, including 2-3 large (> 1 kb) insertions or deletions associated with the movement of mobile DNA elements. Summaries of the genomes and changes observed in strains TP2 and TP3 (relative to TP1) are summarized in **Table 3**, and the locations of AMR genes, transposases, prophages, capsule locus in TP1 genome, and large insertion and deletions (>1 kb) in TP2 and TP3, in reference to TP1, are illustrated in **Figure 4**. In reference to TP1, detailed sequence changes, associated coordinates and genes affected in TP2 and TP3 are listed in **Supplementary Tables S6 and S7**, respectively.

**Figure 4.**
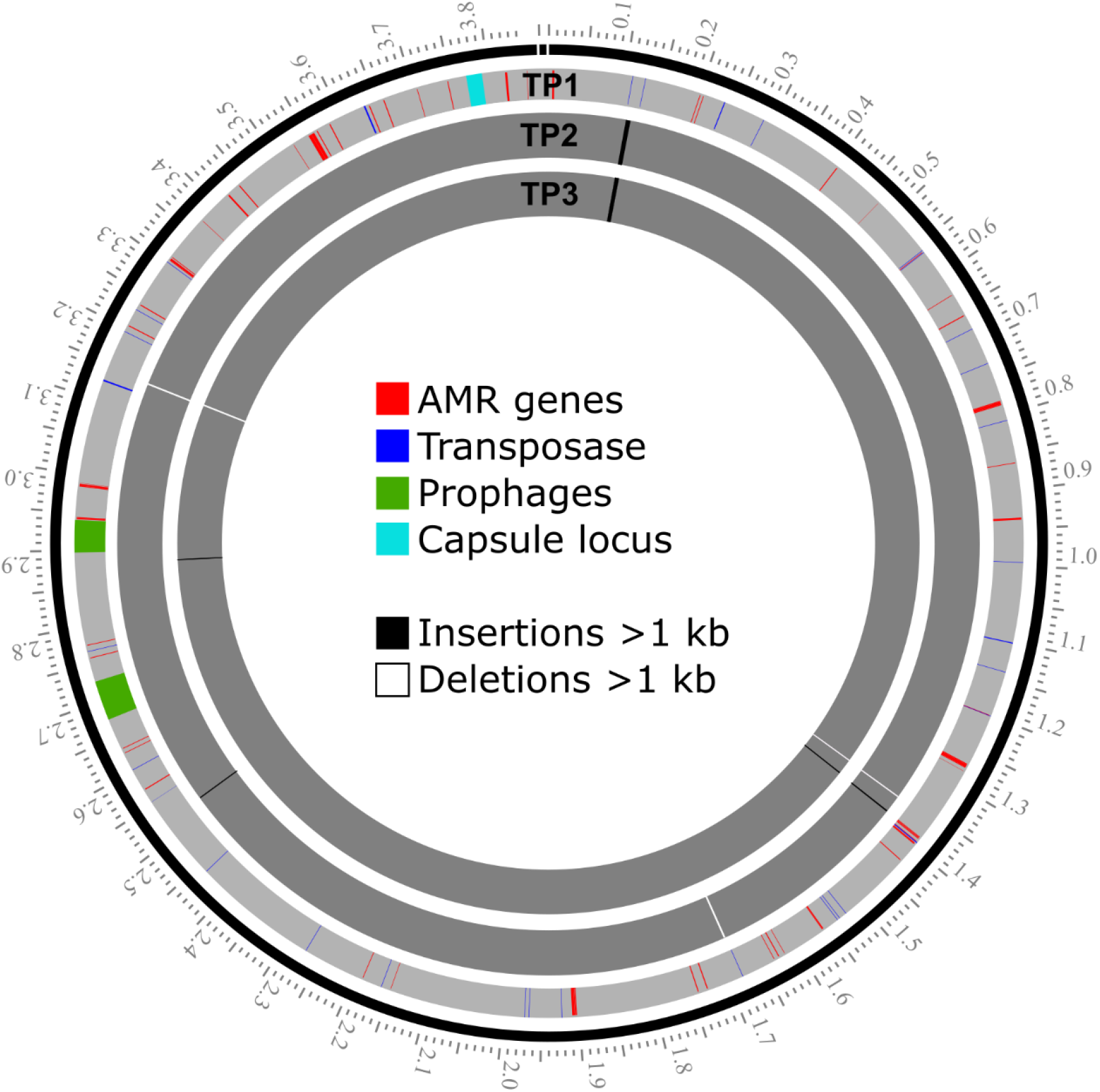
Locations of AMR genes, transposases, prophages, capsule locus in TP1 genome, and large insertion and deletions (>1 kb) in TP2 and TP3, in reference to TP1.

**Figure 5.**
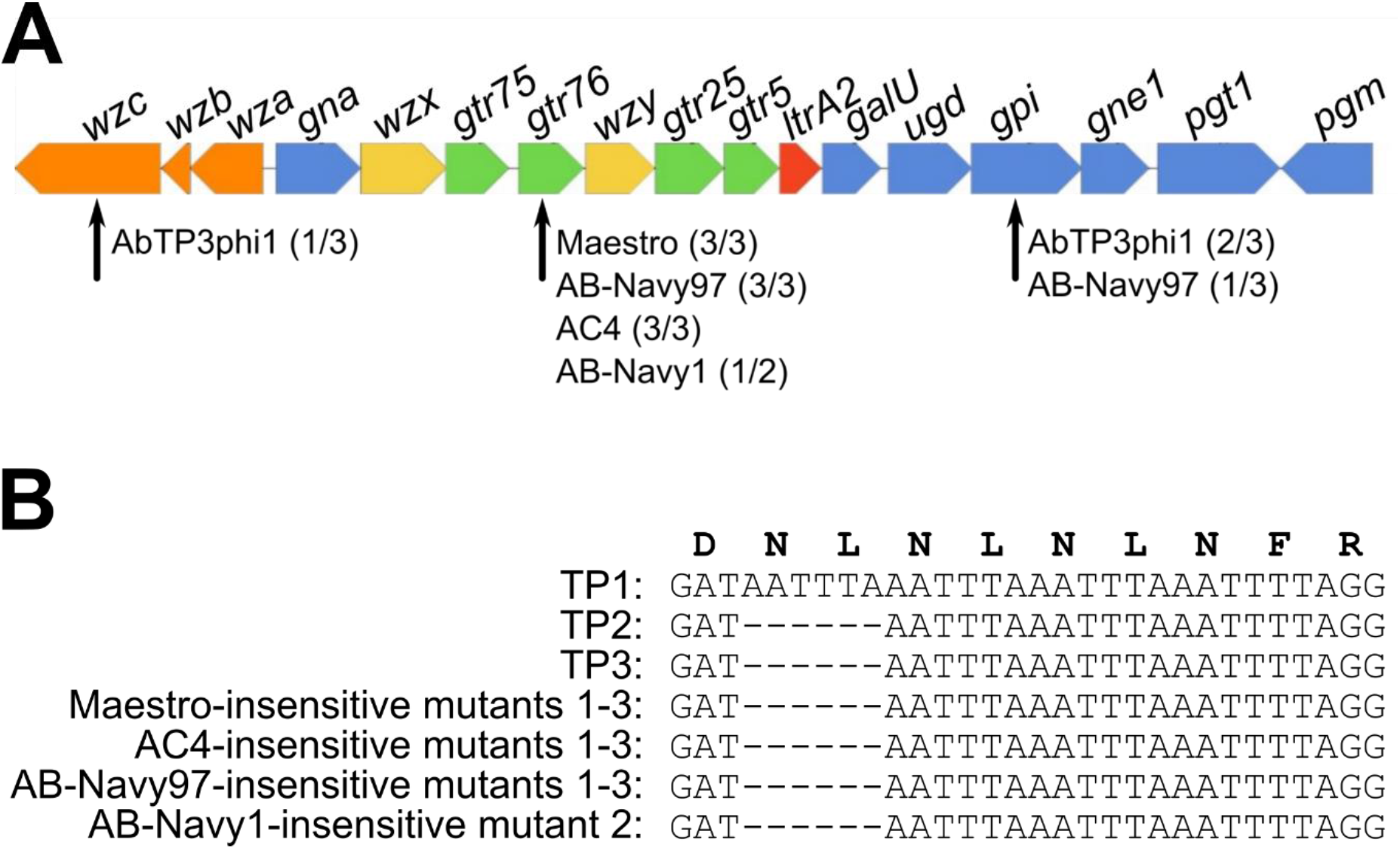
Comparison of the K loci of strains TP1, TP2 and TP3, and phage resistant mutants generated *in vitro*. (A) Nucleotide alignment of the sequences showing the 6 nucleotide deletion in one of the glycosyltransferases (*gtr76*) found in multiple TP1 mutants resistant to the cocktail myophages. (B) Diagram of the KL116 capsule locus identified in strains TP1, TP2, and TP3 as predicted by Kaptive. Genes are represented by arrows oriented in the direction of transcription. Orange arrows represent genes involved in capsule export, yellow genes are involved in repeat unit processing, blue genes are involved in simple sugar biosynthesis, green genes encode glycotransferases and the red gene codes for the initiating transferase. All genes had 100% coverage and ranged from 90% - 100% identity to the KL116 type in the Kaptive database. Defective capsule locus genes identified in *in vitro*-generated phage-insensitive mutants of TP1 are indicated by black arrows; numbers in parentheses after each phage name indicate what proportion of phage-insensitive mutants contained a mutation in that gene.

### Notable large insertions and deletions in TP2 and TP3 compared to TP1

One of the notable changes in TP2 and TP3 is a novel 6,673 bp insertion sequence that is not present in TP1. This element is inserted in a position adjacent to an existing IS3-like transposase at position 111,357 of the TP1 genome (**Figure 4**) (**Supplementary Tables S6, S7**). This insertion introduces a second copy of the same gene such that the new sequence is flanked by identical copies of the transposase, in addition to carrying its own IS30-like transposase. This acquired 6.7 kb element is not native to TP1 and represents an acquisition of new DNA by horizontal gene transfer that occurred during the course of infection, and is most likely the result of DNA acquisition mechanisms unrelated to phage treatment. *A. baumannii* is known for its ability to rapidly acquire mobile DNA elements in the environment via conjugation and natural competence (Wilharm, Piesker et al. 2013, Domingues, Rosario et al. 2019), and to vary surface molecules through horizontal gene transfer (Snitkin, Zelazny et al. 2011). T4-like phages like those used in treatment are generally poor transducers. In phage T4, multiple defects in *ndd*, *denB, 42* and *alc* are required for transduction to occur (Young, Edlin et al. 1982), and these genes are all conserved in the cocktail myophages reported in this study. In addition, transduction requires the phage to be able to productively infect the donor of the acquired DNA, which was likely to have been a different bacterial species and thus insensitive to the phages used.

BLASTn searches of this sequence identified identical or nearly identical sequences in other Gram-negative bacterial genomes or plasmids, including *A. baumannii* (CP038644), *Klebsiella pneumoniae* (LR697132), *E. coli* (CP020524), and *Citrobacter freundii* (KP770032). This inserted sequence encodes a number of significant additional antibiotic resistance determinants, including a predicted aminoglycoside O-phosphotransferase (IPR002575), an NDM-1-like metallo-beta-lactamase (CD16300, IPR001279), and a CutA-like protein that may be involved in metal tolerance (IPR004323). The inserted aminoglycoside O-phosphotransferase (CARD ARO:3003687) is relatively rare in *A. baumannii*, found in 1.43% of *A. baumannii* chromosomes and 0.47% of *A. baumannii* plasmids, as reported by the CARD Resistance Gene Identifier. The prevalence of the inserted NDM-1-like metallo-beta-lactamase (CARD ARO:3000589) is 5.94% of *A. baumannii* genomes and 0.6% of *A. baumannii* plasmids.

Other than the 6.7 kb insertion described above, all other major variations in the TP2 and TP3 genomes can be attributed to recombination, deletion or transposition of elements present in the TP1 genome. Another 1,886 bp insertion sequence was identified in TP2 and TP3, inserted at position 1,381,905 of the TP1 genome, adjacent to an existing IS6-like IS26 family transposase (**Figure 4**) **(Supplementary Tables S6, S7**). This insertion introduces a second copy of the IS6-like transposase and an additional copy of an aminoglycoside O-phosphotransferase (IPR002575) which is also present in TP1 (locus HWQ22_16890) and flanked by copies of the same IS6-like transposase. In this case, this insertion is a duplication of an existing AMR gene rather than the acquisition of foreign DNA. The presence of the new 6.7 kb element and the duplicated 1.9 kb element resulted in 3 extra antibiotic resistance genes in TP2 and TP3 (35 total predicted AMR genes) compared to TP1 (32 total predicted AMR genes) (**Table 3; Supplementary Table S5**).

Strain TP1 contains a 1,094 bp IS30 family transposase present at position 3,143,593 of the chromosome, which is not present in TP2 or TP3. This transposase interrupts a restriction endonuclease-like protein in TP1 which is complete in TP2 and TP3. Based on BLASTn analysis of this region, it appears that the intact state seen in TP2 and TP3 is ancestral, as this version of this region appears in 165 other *Acinetobacter* genomes, while the IS-interrupted version of this region is unique to TP1. This indicates that TP2 and TP3 are not directly descended from TP1. However, TP2 and TP3 do appear to share a common ancestor as they both contain the novel 6.7 kb insertion element, but TP3 is not a clear descendant of TP2 as there are multiple genetic changes in TP2 that are absent in TP3, such as a ~1 kb deletion at the ~1.7 Mb position in TP2 that is not present in TP1 or TP3 (**Figure 4, Supplementary Table S8**). The *A. baumannii* strain was clearly evolving and radiating multiple lineages during the course of infection and treatment. This highlights the fact that bacterial pathogens are undergoing constant selective pressure *in vivo* and do not exist as strictly clonal populations even in a single patient over time.

### Prophage elements in TP1, TP2, and TP3

Prophage analysis revealed two apparently complete prophage regions (52,561 bp and 42,762 bp in length, respectively) in TP1, TP2, and TP3 genomes that are likely to encode active prophages (**Figure 4**). Phage *att* sites and conserved phage proteins (tail and tail tape measure protein, major head subunit and head morphogenesis protein, terminase large subunit, endolysin) were identified in both prophage regions. These two prophage regions are conserved in TP1, TP2, and TP3 and no sequence change was observed among the three strains. The 52 kb prophage is highly conserved (with up to 100% nucleotide identity by BLASTn) in many other *A. baumannii* genomes, including that of ATCC 19606, which is one of the earliest available clinical isolates of *A. baumannii*, dating to the 1940’s (Hamidian, Blasco et al. 2020). This prophage region shares limited similarity to cultured phages, with its closest relative being *Acinetobacter* phage Ab105-3phi (KT588073), with which it shares 49.4% nucleotide identity and 22 similar proteins. The 43 kb prophage region was found to be less conserved in other *A. baumannii* genomes, with the most closely related prophage element sharing only 69% overall sequence identity. This prophage region is ~46% related to *A. baumannii* phage 5W (MT349887), which also appears to be a temperate phage due to the presence of an integrase and LexA-like repressor. Other than 5W, this element is not closely related to any other cultured phages in the NCBI database, sharing no more than 10% nucleotide identity and no more than 8 proteins with other phages.

### Characterization of phage-resistant mutants generated *in vitro* and the comparison to *in vivo* isolates

Five phages selected from the phage cocktails (AC4, Maestro, AB-Navy1, AB-Navy97, AbTP3phi1) were used to select for phage-insensitive mutants *in vitro* using *A. baumannii* strain TP1 as host. Three independent mutants against phages AC4, Maestro, AB-Navy97, AbTP3phi1 were isolated, and two independent mutants against phage AB-Navy1 were isolated. After resequencing and mapping mutant reads to the reference TP1 genome, changes detected with quality scores greater than 100 were examined (**Table 4**). The majority of identified mutations were located in the bacterial KL116 capsule locus. The KL116 capsule is comprised of a five-sugar repeating unit with a three-sugar backbone composed of Gal and GalNAc and a two-sugar side chain composed of Glc and GalNAc (Shashkov, Cahill et al. 2019). In all the mutants resistant to the myophages Maestro, AC4, AB-Navy97, a common 6-bp deletion was observed in a predicted capsular glycosyltransferase protein identified as Gtr76 by Kaptive (HWQ22_04225) (**Figure 5A**). Notably, these 6-bp deletions were also observed in isolates TP2 and TP3, which evolved *in vivo* during phage treatment and were insensitive or showed reduced sensitivity to all myophages tested (Table 2, Supplementary Table S5 and S6). These 6-bp deletions occurred in a region containing four copies of a tandem repeat sequence TAAATT (**Figure 5B**), which probably is prone to mutation by strand slippage events during replication. These mutations result in the deletion of residue L243 and N244, resulting in the loss of a predicted flexible linker between two α-helices in the C-terminus of the glycosyltransferase protein. This protein is predicted to participate in capsule synthesis by forming the β-D-GalNAc-(1 → 4)-D-Gal linkage of the side-chain disaccharide to the trisaccharide backbone (Shashkov, Cahill et al. 2019). Loss of or altered activity in this enzyme would be expected to result in capsule with reduced or absent disaccharide side-chains, suggesting that this side chain plays a major role in host recognition by these phages.

In mutants selected for insensitivity to phage AB-Navy1, one mutant contained the same conserved 6 bp deletion identified in the other mutants, and one lacked this mutation but instead had a nonsense mutation (W183am) in *carO* (HWQ22_09280) (**Table 4**). CarO is a 29 kDa outer membrane transporter, loss of which has been associated with increased antibiotic resistance (Mussi, Limansky et al. 2005, Uppalapati, Sett et al. 2020). The role of CarO in phage sensitivity is not clear, but its truncation may lead to other cell wall defects that reduce sensitivity to this phage; truncations in CarO have been associated with reduced adherence and invasion in tissue culture and with reduced virulence *in vivo* (Labrador-Herrera, Perez-Pulido et al. 2020). This finding illustrates that defects in the capsule locus are not the only means by which TP1 may gain phage insensitivity. Notably, similar CarO defects were not observed in TP2 or TP3, which attained phage resistance *in vivo*.

In addition to the common 6 bp deletion in the Gtr76 glycosyltransferase and CarO mutation, the other mutations observed in the myophage-insensitive mutants are SNPs or small indels in non-coding regions or that result in missense or silent mutations in a predicted capsular glucose-6-phosphate isomerase Gpi (HWQ22_04190) and an ABC transporter, respectively (**Table 4**). However, these SNPs are not conserved in the *in vitro* mutants against myophages and were also not detected in *in vivo* isolates TP2 and TP3, indicating that the defect observed in the Gtr76 glycosyltransferase is sufficient to confer insensitivity to the cocktail myophages in this strain.

Strain TP1 mutants resistant to the podophage AbTP3phi1 were also found to contain mutations in the capsule locus, but these mutations were confined to the genes encoding the glucose-6-phosphate isomerase Gpi and polysaccharide biosynthesis tyrosine autokinase Wzc (HWQ22_04255) (**Table 4, Figure 5A**). Loss of function in these genes is expected to result in loss of L-fructose-6-phosphate required for downstream production of capsule monomers and defects in capsule export, respectively (Wyres, Cahill et al. 2020). This suggests that these mutants may exhibit more severe defects in K116 capsule expression, and that AbTP3phi1 requires the presence of the capsule backbone for successful infection.

Our results are consistent with the recently published work by Altamirano *et al*., where a frameshift in the glycosyltransferase and glucose-6-phosphate isomerase within the K locus were detected in two independent phage-resistant *A. baumannii* mutants, respectively (Gordillo Altamirano, Forsyth et al. 2021). The consistency between our work and that study confirms the *A. baumannii* capsule locus being important for phage sensitivity. Both Gpi and glycosyltransferases are involved the biosynthesis of capsule K units, which are tightly packed repeating subunits consisting of 4 to 6 sugars (Singh, Adams et al. 2018). The reason why one group of phages (our myophages, and the myophage ΦFG02 in Altamirano *et al*.) selected primarily for defects in the Gtr glycosyltransferase but the other phages (our podophage AbTP3phi1 and myophage ΦCO01in Altamirano *et al*.) selected for defects in Gpi is not entirely clear. These phages likely recognize different moieties of the bacterial capsule as their receptors, but it should be noted that many of the mutations associated with insensitivity observed in our study are not necessarily inactivating to the protein: the most common mutation in the capsule locus is a two-residue in-frame deletion in *gtr76* (Figure 4B), and the other mutations are single-residue changes or nonsense/frameshift mutations relatively late in the reading frame. These mutations may modulate protein function rather than being completely inactivating.

Capsule is a known common requirement for *A. baumannii* phages, and defects in capsule synthesis have been shown to be responsible for phage resistance (Billing 1960, Gordillo Altamirano, Forsyth et al. 2021). The presence of the same 6 bp deletion in the capsular glycosyltransferase gene *gtr76* of both the *in vitro-* and the *in vivo*-selected *A. baumannii* strains indicates that the same route to phage insensitivity may be followed by strain TP1 in both systems. Importantly, this demonstrates that laboratory *in vitro* investigations of bacterial selection and phage insensitivity can produce results that are relevant and predictive for the *in vivo* milieu of clinical treatment. However, as can be seen by comparing **Table 3** and **Table 4**, the *in vivo*-selected strains TP2 and TP3 contain numerous genetic changes in addition to those obtained by simple selection *in vitro*. Strains TP2 and TP3, which were recovered from the patient during the course of treatment, evolved in response to nearly continuous antibiotic treatment, to the host immune response to the infection, and to the phage treatment. Thus, many of the genetic changes observed in TP2 and TP3 are likely to be adaptations to these additional stresses, or to compensate for defects in capsule expression in order to survive in this hostile environment.

## Conclusions

In conclusion, this study provides detailed genomic information on the evolution of *A. baumannii* during the course of infection, showing that resistance to the therapeutic phages emerged early, and the acquisition of new mobile elements can occur during treatment. The potential for early emergence of phage resistance should be taken into account when considering the phages to be used for treatment and the optimal duration of the therapeutic regimen. Genomic analysis of the phages used in this intervention illustrates the importance of whole genome sequencing of phages to be used in phage therapy, in addition to the conventional experimental tests for phage host range and growth characteristics. In addition to assessing phage virulence and identifying carriage of potentially deleterious genes, genomic analysis of phage tail fiber proteins is of value in order to select phages utilizing different host recognition mechanisms with a goal of minimizing the development of host resistance, especially considering host receptor identification is a time-consuming experimental process. The use of genetically distinct phages in a phage cocktail can avoid redundancy and significantly save time and effort in phage production and purification, which is also an important consideration in making phage therapy practical. Finally, this work shows that relatively simple *in vitro* selections for host resistance can yield predictive results for how the organism may behave *in vivo* during infection.

## Acknowledgements

The authors would like to thank Drs. Thomas Patterson, Steffanie Strathdee, Sharon Reed at UCSD for their support during this study, Dr. Andrew Hillhouse at Texas A&M Institute for Genome Sciences and Society (TIGSS) for sequencing support. This work was supported by funding from Texas A&M University, Texas AgriLife Research, the National Science Foundation (awards DBI-1565146), and Navy Work Unit Number (WUN) A1417.

Disclaimer: The views expressed in this article are those of the author and do not necessarily reflect the official policy or position of the Department of the Navy, Department of Defense, Department of Veterans Affairs, nor the U.S. Government. The authors Theron Hamilton, Kimberly Bishop-Lilly, and Biswajit Biswas are employees of the U.S. Government, US Government contractor or military service members. This work was prepared as part of official duties. Title 17 U.S.C. §105 provides that “Copyright protection under this title is not available for any work of the United States Government.” Title 17 U.S.C. §101 defines a U.S. Government work as a work prepared by a military service member or employee of the U.S. Government as part of that person’s official duties.

---

**Supplementary Figure 1.**
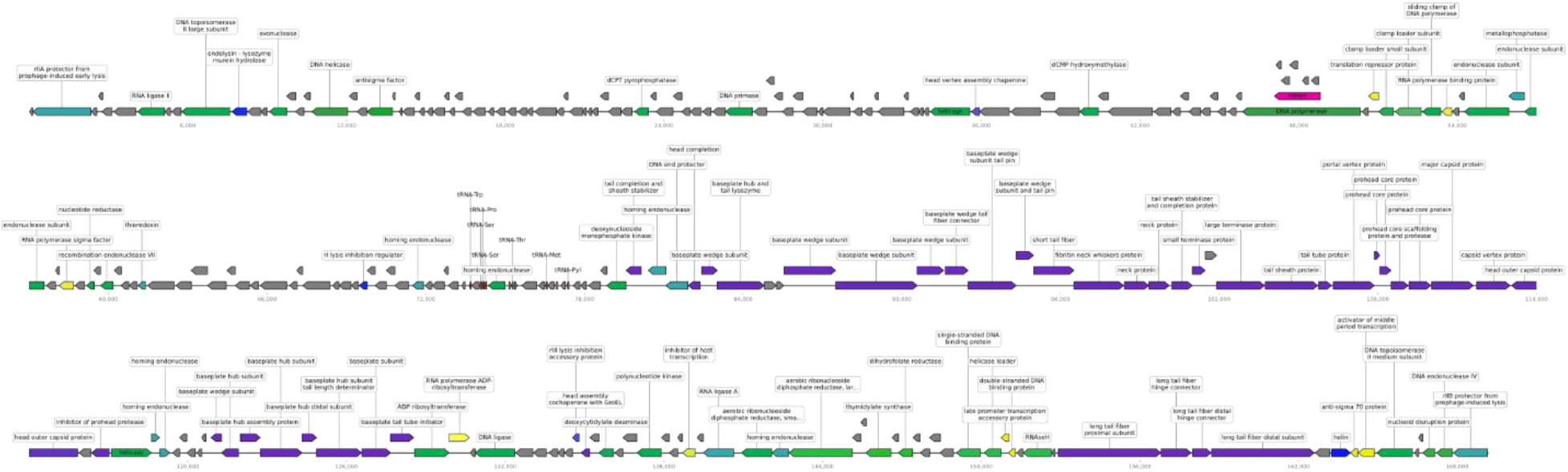
Genomic map of phage Maestro. Predicted genes are represented by blocks. Blocks pointing to the right are genes encoded on the forward strand, pointing to the left are on the reverse strand. The ruler below the genomes indicates scale in bp. Genes are color coded based on functions: regulatory (yellow), DNA replication (green), structural (purple), lysis (blue), intron (pink), tRNA (red), other (turquoise), hypothetical protein with unknown function (grey).

**Supplementary Figure 2.**
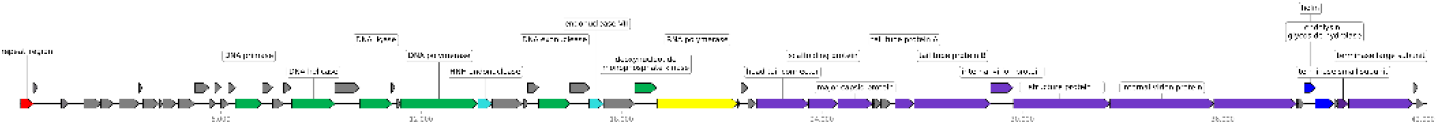
Genome map of AbTP3Phi1. Predicted genes are represented by blocks. Blocks pointing to the right are genes encoded on the forward strand, pointing to the left are on the reverse strand. The ruler below the genomes indicates scale in bp. Genes are color coded based on functions: regulatory (yellow), DNA replication (green), structural (purple), lysis (blue), terminal repeat (red), other (turquoise), hypothetical protein with unknown function (grey).

**Supplementary Figure 3.**
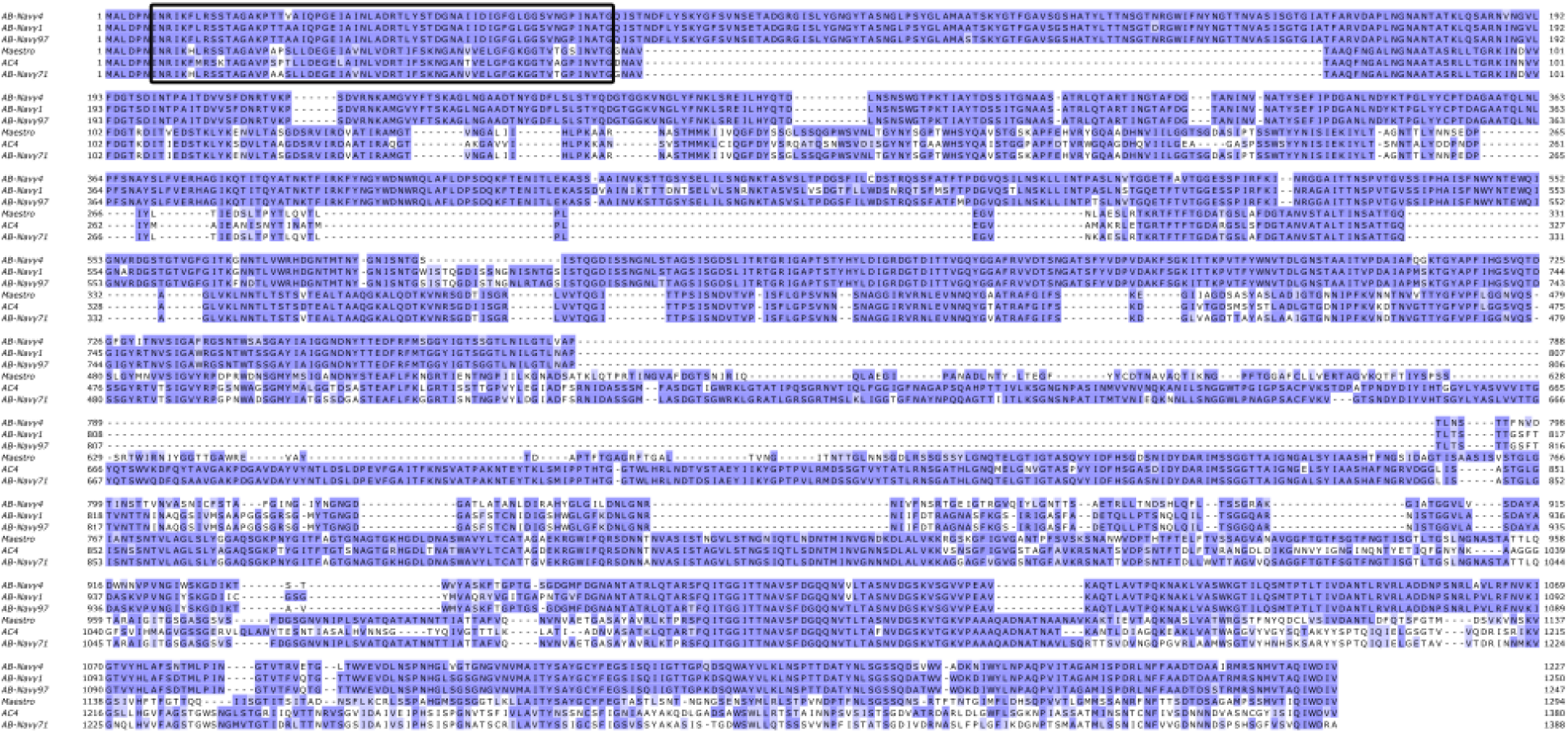
Multiple sequence alignment of the long tail fiber protein sequences of the myophages used in this study. Black box denotes the regions with homology to T4 gp37.

## References

Adams, M. K. (1959). Bactiophages. New York, Interscience Publishers, Inc.

Adriaenssens, E. M., M. Krupovic, P. Knezevic, H. W. Ackermann, J. Barylski, J. R. Brister, M. R. Clokie, S. Duffy, B. E. Dutilh, R. A. Edwards, F. Enault, H. B. Jang, J. Klumpp, A. M. Kropinski, R. Lavigne, M. M. Poranen, D. Prangishvili, J. Rumnieks, M. B. Sullivan, J. Wittmann, H. M. Oksanen, A. Gillis and J. H. Kuhn (2017). “Taxonomy of prokaryotic viruses: 2016 update from the ICTV bacterial and archaeal viruses subcommittee.” Arch Virol 162(4): 1153–1157.

Afgan, E., D. Baker, B. Batut, M. van den Beek, D. Bouvier, M. Cech, J. Chilton, D. Clements, N. Coraor, B. A. Gruning, A. Guerler, J. Hillman-Jackson, S. Hiltemann, V. Jalili, H. Rasche, N. Soranzo, J. Goecks, J. Taylor, A. Nekrutenko and D. Blankenberg (2018). “The Galaxy platform for accessible, reproducible and collaborative biomedical analyses: 2018 update.” Nucleic Acids Res 46(W1): W537–W544.

Alcock, B. P., A. R. Raphenya, T. T. Y. Lau, K. K. Tsang, M. Bouchard, A. Edalatmand, W. Huynh, A. V. Nguyen, A. A. Cheng, S. Liu, S. Y. Min, A. Miroshnichenko, H. K. Tran, R. E. Werfalli, J. A. Nasir, M. Oloni, D. J. Speicher, A. Florescu, B. Singh, M. Faltyn, A. Hernandez-Koutoucheva, A. N. Sharma, E. Bordeleau, A. C. Pawlowski, H. L. Zubyk, D. Dooley, E. Griffiths, F. Maguire, G. L. Winsor, R. G. Beiko, F. S. L. Brinkman, W. W. L. Hsiao, G. V. Domselaar and A. G. McArthur (2020). “CARD 2020: antibiotic resistome surveillance with the comprehensive antibiotic resistance database.” Nucleic Acids Res 48(D1): D517–D525.

Anisimova, M. and O. Gascuel (2006). “Approximate likelihood-ratio test for branches: A fast, accurate, and powerful alternative.” Syst Biol 55(4): 539–552.

Arndt, D., J. R. Grant, A. Marcu, T. Sajed, A. Pon, Y. Liang and D. S. Wishart (2016). “PHASTER: a better, faster version of the PHAST phage search tool.” Nucleic Acids Res 44(W1): W16–21.

Aslam, S., E. Lampley, D. Wooten, M. Karris, C. Benson, S. Strathdee and R. T. Schooley (2020). “Lessons Learned From the First 10 Consecutive Cases of Intravenous Bacteriophage Therapy to Treat Multidrug-Resistant Bacterial Infections at a Single Center in the United States.” Open Forum Infect Dis 7(9): ofaa389.

Bankevich, A., S. Nurk, D. Antipov, A. A. Gurevich, M. Dvorkin, A. S. Kulikov, V. M. Lesin, S. I. Nikolenko, S. Pham, A. D. Prjibelski, A. V. Pyshkin, A. V. Sirotkin, N. Vyahhi, G. Tesler, M. A. Alekseyev and P. A. Pevzner (2012). “SPAdes: a new genome assembly algorithm and its applications to single-cell sequencing.” J Comput Biol 19(5): 455–477.

Bartual, S. G., J. M. Otero, C. Garcia-Doval, A. L. Llamas-Saiz, R. Kahn, G. C. Fox and M. J. van Raaij (2010). “Structure of the bacteriophage T4 long tail fiber receptor-binding tip.” Proc Natl Acad Sci U S A 107(47): 20287–20292.

Billing, E. (1960). “An association between capsulation and phage sensitivity in Erwinia amylovora.” Nature 186: 819–820.

Blankenberg, D., A. Gordon, G. Von Kuster, N. Coraor, J. Taylor, A. Nekrutenko and T. Galaxy (2010). “Manipulation of FASTQ data with Galaxy.” Bioinformatics 26(14): 1783–1785.

Bolger, A. M., M. Lohse and B. Usadel (2014). “Trimmomatic: a flexible trimmer for Illumina sequence data.” Bioinformatics 30(15): 2114–2120.

Cahill, J. and R. Young (2019). “Phage Lysis: Multiple Genes for Multiple Barriers.” Adv Virus Res 103: 33–70.

Camacho, C., G. Coulouris, V. Avagyan, N. Ma, J. Papadopoulos, K. Bealer and T. L. Madden (2009). “BLAST+: architecture and applications.” BMC Bioinformatics 10: 421.

CDC (2019). “ANTIBIOTIC RESISTANCE THREATS IN THE UNITED STATES.” https://www.cdc.gov/drugresistance/biggest-threats.html.

Chevenet, F., C. Brun, A. L. Banuls, B. Jacq and R. Christen (2006). “TreeDyn: towards dynamic graphics and annotations for analyses of trees.” BMC Bioinformatics 7: 439.

Chusri, S., V. Chongsuvivatwong, J. I. Rivera, K. Silpapojakul, K. Singkhamanan, E. McNeil and Y. Doi (2014). “Clinical outcomes of hospital-acquired infection with Acinetobacter nosocomialis and Acinetobacter pittii.” Antimicrob Agents Chemother 58(7): 4172–4179.

Darling, A. E., B. Mau and N. T. Perna (2010). “progressiveMauve: multiple genome alignment with gene gain, loss and rearrangement.” PLoS One 5(6): e11147.

Delcher, A. L., D. Harmon, S. Kasif, O. White and S. L. Salzberg (1999). “Improved microbial gene identification with GLIMMER.” Nucleic Acids Res 27(23): 4636–4641.

Dereeper, A., V. Guignon, G. Blanc, S. Audic, S. Buffet, F. Chevenet, J. F. Dufayard, S. Guindon, V. Lefort, M. Lescot, J. M. Claverie and O. Gascuel (2008). “Phylogeny.fr: robust phylogenetic analysis for the non-specialist.” Nucleic Acids Res 36(Web Server issue): W465–469.

Dijkshoorn, L., A. Nemec and H. Seifert (2007). “An increasing threat in hospitals: multidrug-resistant Acinetobacter baumannii.” Nat Rev Microbiol 5(12): 939–951.

Domingues, S., N. Rosario, A. Candido, D. Neto, K. M. Nielsen and G. J. Da Silva (2019). “Competence for Natural Transformation Is Common among Clinical Strains of Resistant Acinetobacter spp.” Microorganisms 7(2).

Dunn, N. A., D. R. Unni, C. Diesh, M. Munoz-Torres, N. L. Harris, E. Yao, H. Rasche, I. H. Holmes, C. G. Elsik and S. E. Lewis (2019). “Apollo: Democratizing genome annotation.” PLoS Comput Biol 15(2): e1006790.

Edgar, R. C. (2004). “MUSCLE: multiple sequence alignment with high accuracy and high throughput.” Nucleic Acids Res 32(5): 1792–1797.

Garneau, J. R., F. Depardieu, L. C. Fortier, D. Bikard and M. Monot (2017). “PhageTerm: a tool for fast and accurate determination of phage termini and packaging mechanism using next-generation sequencing data.” Sci Rep 7(1): 8292.

Gordillo Altamirano, F., J. H. Forsyth, R. Patwa, X. Kostoulias, M. Trim, D. Subedi, S. K. Archer, F. C. Morris, C. Oliveira, L. Kielty, D. Korneev, M. K. O’Bryan, T. J. Lithgow, A. Y. Peleg and J. J. Barr (2021). “Bacteriophage-resistant Acinetobacter baumannii are resensitized to antimicrobials.” Nat Microbiol 6(2): 157–161.

Gordillo Altamirano, F. L. and J. J. Barr (2019). “Phage Therapy in the Postantibiotic Era.” Clin Microbiol Rev 32(2).

Hamidian, M., L. Blasco, L. N. Tillman, J. To, M. Tomas and G. S. A. Myers (2020). “Analysis of Complete Genome Sequence of Acinetobacter baumannii Strain ATCC 19606 Reveals Novel Mobile Genetic Elements and Novel Prophage.” Microorganisms 8(12).

Harding, C. M., S. W. Hennon and M. F. Feldman (2018). “Uncovering the mechanisms of Acinetobacter baumannii virulence.” Nat Rev Microbiol 16(2): 91–102.

Hernandez-Morales, A. C., L. L. Lessor, T. L. Wood, D. Migl, E. M. Mijalis, J. Cahill, W. K. Russell, R. F. Young and J. J. Gill (2018). “Genomic and Biochemical Characterization of Acinetobacter Podophage Petty Reveals a Novel Lysis Mechanism and Tail-Associated Depolymerase Activity.” J Virol 92(6).

Holt, A., J. Cahill, J. Ramsey, C. Martin, C. O’Leary, R. Moreland, L. T. Maddox, T. Galbadage, R. Sharan, P. Sule, J. D. Cirillo and R. Young (2021). “Phage-encoded cationic antimicrobial peptide required for lysis.” J Bacteriol: JB0021421.

Hyman, P. and M. van Raaij (2018). “Bacteriophage T4 long tail fiber domains.” Biophys Rev 10(2): 463–471.

Jalili, V., E. Afgan, Q. Gu, D. Clements, D. Blankenberg, J. Goecks, J. Taylor and A. Nekrutenko (2020). “The Galaxy platform for accessible, reproducible and collaborative biomedical analyses: 2020 update.” Nucleic Acids Res 48(W1): W395–W402.

Jolley, K. A., J. E. Bray and M. C. J. Maiden (2018). “Open-access bacterial population genomics: BIGSdb software, the PubMLST.org website and their applications.” Wellcome Open Res 3: 124.

Jones, P., D. Binns, H. Y. Chang, M. Fraser, W. Li, C. McAnulla, H. McWilliam, J. Maslen, A. Mitchell, G. Nuka, S. Pesseat, A. F. Quinn, A. Sangrador-Vegas, M. Scheremetjew, S. Y. Yong, R. Lopez and S. Hunter (2014). “InterProScan 5: genome-scale protein function classification.” Bioinformatics 30(9): 1236–1240.

Kongari, R., M. Rajaure, J. Cahill, E. Rasche, E. Mijalis, J. Berry and R. Young (2018). “Phage spanins: diversity, topological dynamics and gene convergence.” BMC Bioinformatics 19(1): 326.

Krieger, I. V., V. Kuznetsov, J. Y. Chang, J. Zhang, S. H. Moussa, R. F. Young and J. C. Sacchettini (2020). “The Structural Basis of T4 Phage Lysis Control: DNA as the Signal for Lysis Inhibition.” J Mol Biol 432(16): 4623–4636.

Krogh, A., B. Larsson, G. von Heijne and E. L. Sonnhammer (2001). “Predicting transmembrane protein topology with a hidden Markov model: application to complete genomes.” J Mol Biol 305(3): 567–580.

Labrador-Herrera, G., A. J. Perez-Pulido, R. Alvarez-Marin, C. S. Casimiro-Soriguer, T. Cebrero-Cangueiro, J. Moran-Barrio, J. Pachon, A. M. Viale and M. E. Pachon-Ibanez (2020). “Virulence role of the outer membrane protein CarO in carbapenem-resistant Acinetobacter baumannii.” Virulence 11(1): 1727–1737.

Langmead, B. and S. L. Salzberg (2012). “Fast gapped-read alignment with Bowtie 2.” Nat Methods 9(4): 357–359.

Langmead, B., C. Trapnell, M. Pop and S. L. Salzberg (2009). “Ultrafast and memory-efficient alignment of short DNA sequences to the human genome.” Genome Biol 10(3): R25.

Laslett, D. and B. Canback (2004). “ARAGORN, a program to detect tRNA genes and tmRNA genes in nucleotide sequences.” Nucleic Acids Res 32(1): 11–16.

Lee, C. R., J. H. Lee, M. Park, K. S. Park, I. K. Bae, Y. B. Kim, C. J. Cha, B. C. Jeong and S. H. Lee (2017). “Biology of Acinetobacter baumannii: Pathogenesis, Antibiotic Resistance Mechanisms, and Prospective Treatment Options.” Front Cell Infect Microbiol 7: 55.

Lee, I. M., I. F. Tu, F. L. Yang, T. P. Ko, J. H. Liao, N. T. Lin, C. Y. Wu, C. T. Ren, A. H. Wang, C. M. Chang, K. F. Huang and S. H. Wu (2017). “Structural basis for fragmenting the exopolysaccharide of Acinetobacter baumannii by bacteriophage PhiAB6 tailspike protein.” Sci Rep 7: 42711.

Madeira, F., Y. M. Park, J. Lee, N. Buso, T. Gur, N. Madhusoodanan, P. Basutkar, A. R. N. Tivey, S. C. Potter, R. D. Finn and R. Lopez (2019). “The EMBL-EBI search and sequence analysis tools APIs in 2019.” Nucleic Acids Res 47(W1): W636–W641.

Miller, E. S., E. Kutter, G. Mosig, F. Arisaka, T. Kunisawa and W. Ruger (2003). “Bacteriophage T4 genome.” Microbiol Mol Biol Rev 67(1): 86–156, table of contents.

Mussi, M. A., A. S. Limansky and A. M. Viale (2005). “Acquisition of resistance to carbapenems in multidrug-resistant clinical strains of Acinetobacter baumannii: natural insertional inactivation of a gene encoding a member of a novel family of beta-barrel outer membrane proteins.” Antimicrob Agents Chemother 49(4): 1432–1440.

Nobrega, F. L., M. Vlot, P. A. de Jonge, L. L. Dreesens, H. J. E. Beaumont, R. Lavigne, B. E. Dutilh and S. J. J. Brouns (2018). “Targeting mechanisms of tailed bacteriophages.” Nat Rev Microbiol 16(12): 760–773.

Noguchi, H., T. Taniguchi and T. Itoh (2008). “MetaGeneAnnotator: detecting species-specific patterns of ribosomal binding site for precise gene prediction in anonymous prokaryotic and phage genomes.” DNA Res 15(6): 387–396.

Peleg, A. Y., H. Seifert and D. L. Paterson (2008). “Acinetobacter baumannii: emergence of a successful pathogen.” Clin Microbiol Rev 21(3): 538–582.

Popova, A. V., D. G. Lavysh, E. I. Klimuk, M. V. Edelstein, A. G. Bogun, M. M. Shneider, A. E. Goncharov, S. V. Leonov and K. V. Severinov (2017). “Novel Fri1-like Viruses Infecting Acinetobacter baumannii-vB_AbaP_AS11 and vB_AbaP_AS12-Characterization, Comparative Genomic Analysis, and Host-Recognition Strategy.” Viruses 9(7).

Ramsey, J., H. Rasche, C. Maughmer, A. Criscione, E. Mijalis, M. Liu, J. C. Hu, R. Young and J. J. Gill (2020). “Galaxy and Apollo as a biologist-friendly interface for high-quality cooperative phage genome annotation.” PLoS Comput Biol 16(11): e1008214.

Regeimbal, J. M., A. C. Jacobs, B. W. Corey, M. S. Henry, M. G. Thompson, R. L. Pavlicek, J. Quinones, R. M. Hannah, M. Ghebremedhin, N. J. Crane, D. V. Zurawski, N. C. Teneza-Mora, B. Biswas and E. R. Hall (2016). “Personalized Therapeutic Cocktail of Wild Environmental Phages Rescues Mice from Acinetobacter baumannii Wound Infections.” Antimicrob Agents Chemother 60(10): 5806–5816.

Roca, I., P. Espinal, X. Vila-Farres and J. Vila (2012). “The Acinetobacter baumannii Oxymoron: Commensal Hospital Dweller Turned Pan-Drug-Resistant Menace.” Front Microbiol 3: 148.

Schooley, R. T., B. Biswas, J. J. Gill, A. Hernandez-Morales, J. Lancaster, L. Lessor, J. J. Barr, S. L. Reed, F. Rohwer, S. Benler, A. M. Segall, R. Taplitz, D. M. Smith, K. Kerr, M. Kumaraswamy, V. Nizet, L. Lin, M. D. McCauley, S. A. Strathdee, C. A. Benson, R. K. Pope, B. M. Leroux, A. C. Picel, A. J. Mateczun, K. E. Cilwa, J. M. Regeimbal, L. A. Estrella, D. M. Wolfe, M. S. Henry, J. Quinones, S. Salka, K. A. Bishop-Lilly, R. Young and T. Hamilton (2017). “Development and Use of Personalized Bacteriophage-Based Therapeutic Cocktails To Treat a Patient with a Disseminated Resistant Acinetobacter baumannii Infection.” Antimicrob Agents Chemother 61(10).

Shashkov, A. S., S. M. Cahill, N. P. Arbatsky, A. C. Westacott, A. A. Kasimova, M. M. Shneider, A. V. Popova, D. A. Shagin, A. A. Shelenkov, Y. V. Mikhailova, Y. G. Yanushevich, M. V. Edelstein, J. J. Kenyon and Y. A. Knirel (2019). “Acinetobacter baumannii K116 capsular polysaccharide structure is a hybrid of the K14 and revised K37 structures.” Carbohydr Res 484: 107774.

Singh, J. K., F. G. Adams and M. H. Brown (2018). “Diversity and Function of Capsular Polysaccharide in Acinetobacter baumannii.” Front Microbiol 9: 3301.

Snitkin, E. S., A. M. Zelazny, C. I. Montero, F. Stock, L. Mijares, N. C. S. Program, P. R. Murray and J. A. Segre (2011). “Genome-wide recombination drives diversification of epidemic strains of Acinetobacter baumannii.” Proc Natl Acad Sci U S A 108(33): 13758–13763.

Summer, E. J. (2009). “Preparation of a phage DNA fragment library for whole genome shotgun sequencing.” Methods Mol Biol 502: 27–46.

Tatusova, T., M. DiCuccio, A. Badretdin, V. Chetvernin, E. P. Nawrocki, L. Zaslavsky, A. Lomsadze, K. D. Pruitt, M. Borodovsky and J. Ostell (2016). “NCBI prokaryotic genome annotation pipeline.” Nucleic Acids Res 44(14): 6614–6624.

UniProt Consortium, T. (2018). “UniProt: the universal protein knowledgebase.” Nucleic Acids Res 46(5): 2699.

Uppalapati, S. R., A. Sett and R. Pathania (2020). “The Outer Membrane Proteins OmpA, CarO, and OprD of Acinetobacter baumannii Confer a Two-Pronged Defense in Facilitating Its Success as a Potent Human Pathogen.” Front Microbiol 11: 589234.

Vaas, L. A., J. Sikorski, B. Hofner, A. Fiebig, N. Buddruhs, H. P. Klenk and M. Goker (2013). “opm: an R package for analysing OmniLog(R) phenotype microarray data.” Bioinformatics 29(14): 1823–1824.

Walker, B. J., T. Abeel, T. Shea, M. Priest, A. Abouelliel, S. Sakthikumar, C. A. Cuomo, Q. Zeng, J. Wortman, S. K. Young and A. M. Earl (2014). “Pilon: an integrated tool for comprehensive microbial variant detection and genome assembly improvement.” PLoS One 9(11): e112963.

Wick, R. R., E. Heinz, K. E. Holt and K. L. Wyres (2018). “Kaptive Web: User-Friendly Capsule and Lipopolysaccharide Serotype Prediction for Klebsiella Genomes.” J Clin Microbiol 56(6).

Wilharm, G., J. Piesker, M. Laue and E. Skiebe (2013). “DNA uptake by the nosocomial pathogen Acinetobacter baumannii occurs during movement along wet surfaces.” J Bacteriol 195(18): 4146–4153.

Wyres, K. L., S. M. Cahill, K. E. Holt, R. M. Hall and J. J. Kenyon (2020). “Identification of Acinetobacter baumannii loci for capsular polysaccharide (KL) and lipooligosaccharide outer core (OCL) synthesis in genome assemblies using curated reference databases compatible with Kaptive.” Microb Genom 6(3).

Young, K. K., G. J. Edlin and G. G. Wilson (1982). “Genetic analysis of bacteriophage T4 transducing bacteriophages.” J Virol 41(1): 345–347.

Young, R. and J. J. Gill (2015). “MICROBIOLOGY. Phage therapy redux--What is to be done?” Science 350(6265): 1163–1164.

